# Distinct Spatiotemporal Representations of Image Beauty and Image Quality

**DOI:** 10.64898/2026.08.20.745789

**Authors:** Philipp Flieger, Rico Stecher, Daniel Kaiser

## Abstract

Humans rapidly assess the beauty of natural scene images. Previous EEG work suggests neural representations of beauty emerge early and are temporally sustained. Complementary fMRI work pinpoints the neural correlates of beauty to visual, frontal, and default-mode network areas. An integrated view of the spatiotemporal dynamics that give rise to the perception of beauty, however, is lacking. Beyond the beauty of the depicted scene, the quality of the image itself influences its perceived beauty, and it is unknown how the brain separates these two factors. To address these questions, we recorded EEG (*N* = 52) and fMRI (*N* = 29) data while participants rated the beauty of 100 natural scene photographs. Another group of participants (*N* = 46) rated the image quality of the same photographs. Separate representational similarity analyses on the EEG and fMRI data revealed early and sustained beauty-related representations across widespread cortical areas. In contrast, representations of image quality emerged earlier, had markedly different representational dynamics, and were predominantly localized to visual cortex. In a model-based EEG-fMRI fusion analysis, we investigated how the correspondence between temporally resolved EEG signals and spatially resolved fMRI signals is explained by beauty ratings. Our results suggest that beauty-related representations emerge early (from around 165ms and peaking at 275ms post-onset), are long-lasting, and primarily originate from high-level visual cortex. This spatiotemporal signature persisted when controlling for image-quality ratings. Our findings emphasize the importance of perceptual processing for perceived beauty and suggest that the brain represents aesthetic appeal independently of image quality.

## Introduction

Humans are attracted by visual beauty, and we actively seek beautiful experiences in museums or cinemas. Yet, aesthetic experiences are common not only in places like museums but we spontaneously encounter them in daily life, too, such as when we see a beautiful sunset or colorful fireworks (cf. Brown & Dissanayake, 2018). How does the brain represent the beauty of such natural stimuli? Neuroimaging work has revealed that the assessment of visual beauty is mediated by a complex network of brain regions associated with perception, executive control, reward, and self-referential processing (Chatterjee & Vartanian, 2014; Iigaya et al., 2023; Nadal & Skov, 2024; Starr, 2023; Vessel et al., 2019). Fewer studies have investigated the temporal dynamics of beauty-related representations. Evidence from M/EEG experiments suggests that representations of beauty emerge during perceptual processing (as early as 200ms) and are relatively sustained over time (Kaiser, 2022a, 2022b; Kaiser & Nyga, 2020; Lin et al., 2024; Muñoz & Martín-Loeches, 2015; Strijbosch et al., 2022).

Most theories of neuroaesthetics agree that perceived beauty is already, at least in part, resolved during perceptual processing (Chatterjee & Vartanian, 2014; Karim et al., 2022; Pearce et al., 2016). This perceptual basis of beauty is linked to early responses in the M/EEG (Kaiser, 2022a, 2022b; Kaiser & Nyga, 2020; Muñoz & Martín-Loeches, 2015) that presumably correspond to fMRI activations in visual cortex (Isik & Vessel, 2021). In line with this idea, deep neural network models of vision predict perceived beauty well, even when their training objectives are orthogonal to beauty judgments (Conwell et al., 2025; Damiano et al., 2023; Nara & Kaiser, 2024).

By and large, these previous studies investigated beauty by using images of the real world. Yet, the beauty of an image depends not only on the beauty of what is depicted, but also on the quality of the image. The influence of image quality and how it interacts with the beauty of the depicted content are relatively underexplored (Tinio & Leder, 2009). Prior research demonstrated that more degraded (in terms of contrast, sharpness, and grain) photographs are liked less than higher-fidelity versions of the same stimuli (Tinio et al., 2011). This suggests a systematic difference between the perception of beauty in the outside world (e.g., looking at a landscape during a hike) compared to the perception of beauty in depictions of the world (e.g., looking at landscape photography). However, it is currently unclear how the perceived quality of a stimulus influences neural representations of its aesthetic appeal. Specifically, we do not know how the brain represents the beauty and image quality of a stimulus independently from each other, and how representations of beauty and image quality evolve differently across time and space.

Beyond perception, beauty judgments are mediated by a set of cognitive processes linked to activations in higher-order brain regions. First, it has been suggested that the default-mode network (DMN) is critically involved in the representation of beauty (e.g., Belfi et al., 2019; Vessel et al., 2012, 2013) and codes beauty in ways that generalize across categories (Vessel et al., 2019). Second, higher-order attentional processes likely to contribute to the representations of beauty as they help identify and maintain task and stimulus attributes for further processing (e.g., focusing on the façade of a building versus judging it in the context of the surrounding environment; Cattaneo, 2020; Cela-Conde et al., 2013; Grosbras et al., 2012). Finally, beauty judgments require the interpretation of valuation and reward signals from the medial prefrontal (PFCm) and orbitofrontal (OFC) cortices. Taken together, the neural coding of beauty thus comprises a perceptual stage, during which perceptual stimulus attributes are evaluated, and a cognitive stage, during which self-referential processing, attention, and reward signals shape beauty judgments.

There are several open questions about how representations of image beauty and image quality evolve throughout the course of this processing cascade. First, how do neural representations of beauty emerge concurrently across space and time? Second, does the human brain process image quality independently of beauty? Third, how do representations of beauty and image quality differ across processing time and brain space? To answer these questions, we obtained behavioral beauty ratings of 100 natural scene images in separate experiments while participants’ brain activity was recorded using EEG (*N* = 52) and fMRI (*N* = 29). We additionally collected image-quality ratings for the same stimuli in an independent survey (*N* = 46). Performing representational similarity analysis (RSA; Kriegeskorte et al., 2008) on the individual fMRI and EEG datasets allowed us to characterize the spatial topography and temporal dynamics of beauty and image-quality representations, respectively. Our fMRI results confirm previous findings for widespread beauty representations all over the cortex (Iigaya et al., 2023; Vessel et al., 2019), while our EEG results substantiate that these representations emerge early and are sustained over time (Kaiser, 2022a; Kaiser & Nyga, 2020; Nara et al., 2026). Image-quality representations, in comparison, were predominantly limited to visual cortex and emerged more rapidly than beauty representations. Critically, beauty-related representations persisted when controlling for image quality, suggesting that beauty and image quality are processed independently of one another. Furthermore, we characterized the neural correlates of beauty in a spatiotemporally resolved analysis, combining EEG and fMRI data using a model-based fusion approach (Cichy et al., 2014; Cichy & Oliva, 2020). Performing this analysis across a set of 50 brain parcels yielded a time course of beauty-related representations for each parcel. We found that beauty representations in the EEG signals primarily originated from three parcels in ventral visual cortex, suggesting that perceived scene beauty is most prominently related to high-level visual representations.

## Methods

### Participants

#### fMRI

Thirty healthy adults (*µ_age_* = 27.7 years, *SD_age_* = 4.8 years, 18 female) signed up via a recruitment email disseminated among students and employees of Justus Liebig University Giessen. One participant was excluded due to excessive head motion, yielding a final sample of 29 participants. All participants were right-handed, had normal or corrected-to-normal vision, and no history of psychiatric or neurological disorders. Participants provided written consent prior to their participation and received a financial reward for partaking in the study. All procedures were approved by the ethics committee of Justus Liebig University Giessen and were in accordance with the Declaration of Helsinki.

#### EEG

We used EEG recordings from a total of 52 EEG participants that were tested in two previous experiments (Kaiser, 2022a; Nara et al., 2026). Twenty-four healthy adults (*M_age_* = 19.6 years, *SD_age_* = 1.7 years, 21 female) took part in the first EEG experiment (Kaiser, 2022a). One participant had to be excluded due to a recording error, yielding a final sample of 23 participants. Procedures were approved by the ethics committees of the Department of Psychology, University of York. Another 30 healthy adults (*M_age_* = 24.5 years, *SD_age_* = 4.1 years, 21 female) took part in the second EEG experiment (Nara et al., 2026). One participant had to be excluded due to excessive head movement, yielding a final sample of 29 participants. Procedures were approved by the ethics committees of the Justus Liebig University Giessen. In both studies, participants provided written informed consent prior to their participation. Both experiments were in accordance with the Declaration of Helsinki. Participants received course credits or monetary compensation for their participation.

### Stimuli

We used the stimulus set from Kaiser (2022a) in both the EEG and fMRI experiments. The stimuli consisted of 100 diverse, natural scene photographs, resized to 600×400 pixels. The images were taken from the AVA (Murray et al., 2012) and photo.net (Datta et al., 2008) aesthetics-rating databases. Stimuli were arranged in 50 pairs. In each pair, one stimulus was highly rated for aesthetic appeal and one was poorly rated (based on the databases). Pairs were matched in terms of their overall content (e.g., whether a natural scene or human was depicted) and low-level visual features (Figure 1A; for details see: Kaiser, 2022a).

**Figure 1.**
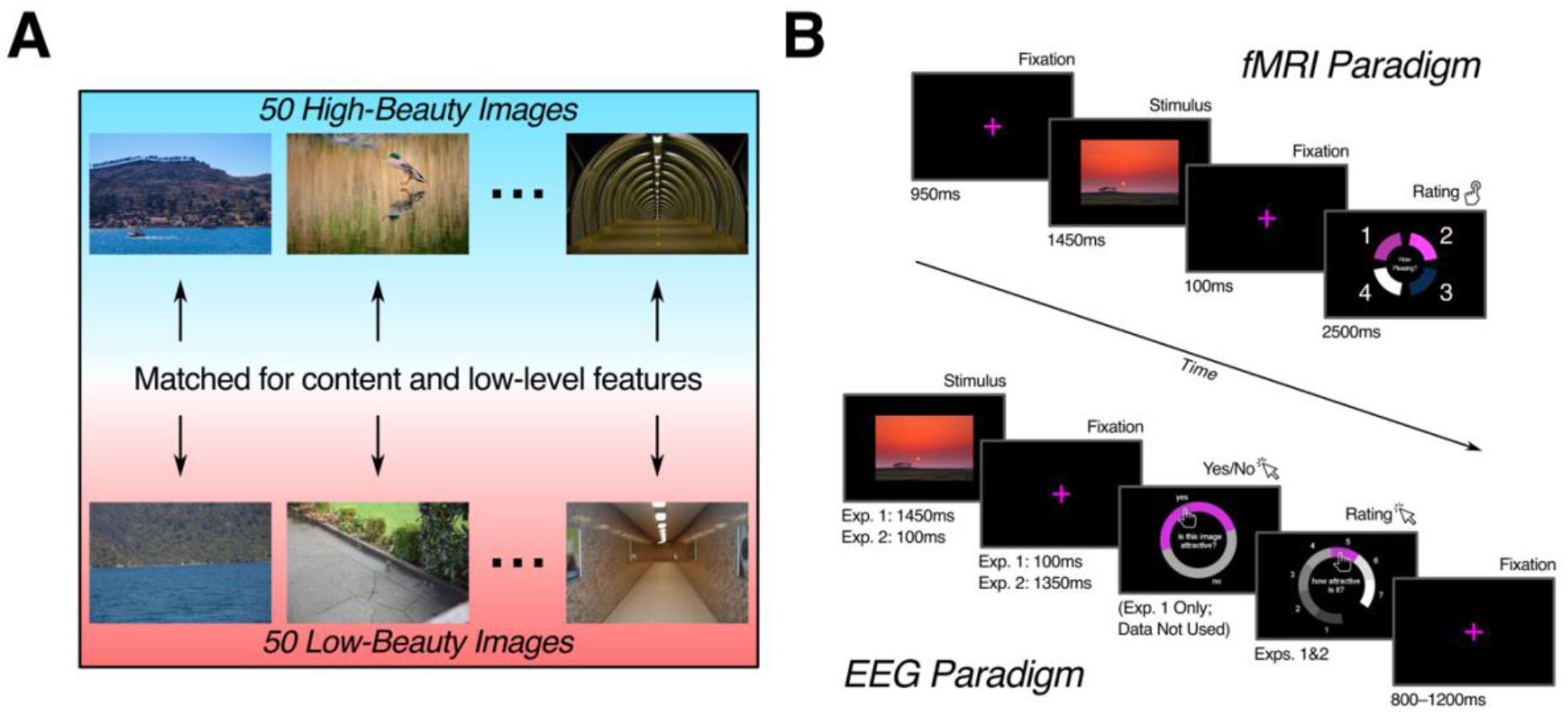
Overview of stimuli and experimental paradigms. *A*. Example stimulus pairs with high- and low-rated counterparts. During each run in both the fMRI and EEG experiments, participants viewed 100 natural scenes, divided into 50 pairs with one rated as more, and the other as less beautiful (based on database ratings; see Methods: Stimuli for details). *B.* Neuroimaging paradigms. During fMRI (top), a single scene was shown on each trial followed by a rating response made via button press (buttons corresponding to wedges on circular response wheel; 1: least beautiful, 4: most beautiful). We used EEG data from two previous EEG studies with very similar designs (bottom; experiment 1: (Kaiser, 2022a); experiment 2: (Nara et al., 2026); see Methods: Paradigm for details). Participants in experiment 1 were asked to give a binary response to indicate whether the previously shown stimulus was attractive to them or not, however, these yes/no responses were not included in the current study. We only used the continuous beauty ratings (1: least beautiful, 7: most beautiful) made via mouse click on a circular response wheel.

### Paradigm

#### fMRI

Stimulus presentation and response collection was controlled using Psychtoolbox (Brainard, 1997) for MATLAB. fMRI participants performed a slightly modified version of the previous EEG experiments (see below). Each trial started with a 950ms fixation period before the stimulus was shown for 1450ms. After another 100ms interval, participants provided their beauty-rating response. Here, the stimuli were rated on a scale from one to four (one = least attractive, four = most attractive), matching the button configuration in our fMRI response device. Each rating was located next to a curved wedge arranged around four quadrants of a circular response screen. The position of the wedges was randomized on each trial to prevent the participant from preparing a motor response (ratings always increased going clockwise). Participants were given 2500ms to provide their ratings. Every stimulus was shown once in each of the seven scanning runs, adding up to seven repetitions per stimulus and a total of 700 trials.

#### EEG

We combined EEG data from two previous EEG experiments (Kaiser, 2022a; Nara et al., 2026) that used the same stimuli to increase power and sample size. Even though stimulus-presentation times differed in the studies, the RSA correlation time courses were highly similar (see: Nara et al., 2026). In both studies, stimulus presentation and response collection was controlled using Psychtoolbox (Brainard, 1997) for MATLAB. In each trial of the first EEG experiment (Kaiser, 2022a), a single scene was shown for 1450ms, followed by a 100ms blank screen, and finally two subsequent rating prompts. First, the participant gave a simple yes/no response as to whether they found the scene attractive or not. Second, they had to indicate how attractive they found the scene on a scale from one to seven (one = least attractive, seven = most attractive). Responses were provided with the computer mouse. To prevent participants from preparing a motor response, the response options for both the yes/no and the rating-scale prompts were located at random angular locations on a circular response screen (analogous setup to fMRI experiment). For the purpose of the current study, only the rating responses were used. Trials were separated by an inter-trial interval randomly varying between 800 and 1200ms. Similar to the fMRI study, there were seven blocks in total, during each of which every image was shown once, yielding seven repetitions per scene and a total of 700 trials. The second EEG experiment (Nara et al., 2026) was identical to the first experiment, with two exceptions. First, scenes were shown for 100ms only, followed by 1350ms of fixation prior to the response. Second, participants only provided the one-to-seven rating response on each trial (i.e., the yes/no response was omitted).

### Neuroimaging Data Acquisition and Preprocessing

#### fMRI

The MRI data were acquired using a 3T Siemens Magnetom PRISMA scanner (Siemens, Erlangen, Germany) and a 64-channel head coil. T_2_*-weighted functional images were collected in descending order using a gradient-echo EPI sequence (TR = 1850ms, TE = 30ms, 75° flip angle, 2.2mm^3^ voxel size, 58 slices, 20% gap/distance factor, 220mm FOV, 100×100 matrix size, descending acquisition). T_1_-weighted anatomical images were acquired using MPRAGE (1mm^3^ voxel size). The fMRI data were preprocessed using SPM12 (www.fil.ion.ucl.ac.uk/spm) and custom MATLAB code. The functional volumes were first realigned and coregistered to the T_1_ image. Afterward both the anatomical and functional images were normalized to Montreal Neurological Institute (MNI) standard space. Finally, we used GLMsingle (Prince et al., 2022) to model the fMRI response to each stimulus during each run, including 3D rotation and translation as motion regressors. This involved selecting a custom hemodynamic response function for each voxel, data denoising using cross-validated principal component analysis, and a voxel-wise regularization of beta estimates via fractional ridge regression. The resulting beta maps were used for both the RSA and EEG-fMRI fusion.

#### fMRI Parcellation

We conducted our analyses in 50 *a priori* defined brain parcels from a cortical parcellation based on resting-state functional connectivity (Yan et al., 2023). All parcel masks were normalized to MNI standard space. Given that the parcels in each hemisphere are homotopic to their contralateral counterparts (i.e., each parcel’s voxels are in approximately the same location in both hemispheres), we created a mask that featured both the left and right aspect of each parcel, yielding a total of 50 bilateral parcels. Moreover, since the parcels are organized according to the functional networks they are part of (e.g., DMN, limbic, somatomotor; Yan et al., 2023), we rearranged the parcels accordingly in our analyses.

#### EEG

In the first EEG experiment (Kaiser, 2022a), EEG data were recorded using an ANT Waveguard 64-channel electrode system and a TMSi REFA amplifier. In the second experiment (Nara et al., 2026) data were recorded using a 64-channel BrainVision recorder with an Actichamp amplifier. Except for the different recording systems, the preprocessing steps were identical. Offline preprocessing was performed in FieldTrip (Oostenveld et al., 2011). The EEG data were referenced to the Fz electrode online and were initially epoched from -500ms to 1900ms relative to stimulus onset. Channels and trials featuring excessive noise were removed based on visual inspection, eye blink artefacts were removed using independent component analysis. After preprocessing, a time window from -250ms to 1450ms relative to stimulus onset was chosen for further analysis.

### Measuring Neural Representational Similarity

To track neural representations in space and time, we performed RSA (Kriegeskorte et al., 2008) on both the fMRI and EEG data using CoSMoMVPA (Oosterhof et al., 2016) for MATLAB.

#### fMRI

We extracted representational dissimilarity matrices (RDMs) from every set of voxels, that is, either the voxels falling within searchlight spheres centered on every voxel (for the RSA) or the voxels falling within each of the 50 parcels (for the RSA and fusion analysis). For each participant and in every set of voxels, we then averaged the beta values across all stimulus presentations, correlated (Spearman correlation) the response vectors for each pair of stimuli and subtracted the correlations from one, yielding a single 100×100 RDM per participant for the given set of voxels (i.e., for each searchlight sphere or parcel).

#### EEG

We extracted response patterns across electrodes for consecutive bins of 10ms each. The data at each bin were unfolded into a vector, on which we performed principal component analysis (PCA) to reduce their dimensionality (Grootswagers et al., 2017; Kaiser et al., 2020). The resulting data were split into two independent subsets, each of which was randomly assigned an equal number of trials per scene. We performed the PCA decomposition on the first subset, which was projected onto the second subset. We only retained the components needed to explain 99% of the variance. Afterward, EEG RDMs were created based on the second subset by averaging across all trials for each scene, calculating the correlation distance (here: 1 - Spearman correlation) between the neural response vectors of each pair of scenes and then reshaping the pairwise distances into a 100×100 RDM with each element now indexing the neural dissimilarity between every pair of scenes. The procedure was repeated with the two subsets permuted. The entire analysis was repeated for a total of 50 times, with random assignment of the trials to one of the subsets each time. Finally, the RDMs were averaged across all repetitions, yielding a single RDM for each bin.

### Modeling Representational Similarity

#### Beauty Ratings

We constructed a beauty predictor RDM using all the behavioral beauty ratings obtained in the EEG and fMRI studies. To this end, we first calculated the average beauty rating for each stimulus across subjects and runs of the fMRI study. Afterward, we computed the pairwise rating differences across all stimuli obtained in the fMRI study. Because we used a four-point instead of seven-point rating scale in the scanner, we rescaled the pairwise rating differences to a seven-point system using linear interpolation. The same procedure was repeated for the EEG, without interpolation. The beauty-rating RDMs from the fMRI and EEG studies (*r* = 0.88) as well as from the two EEG studies (*r* = 0.93) were highly correlated, so we averaged them into a single beauty predictor RDM (Figure 2A).

**Figure 2.**
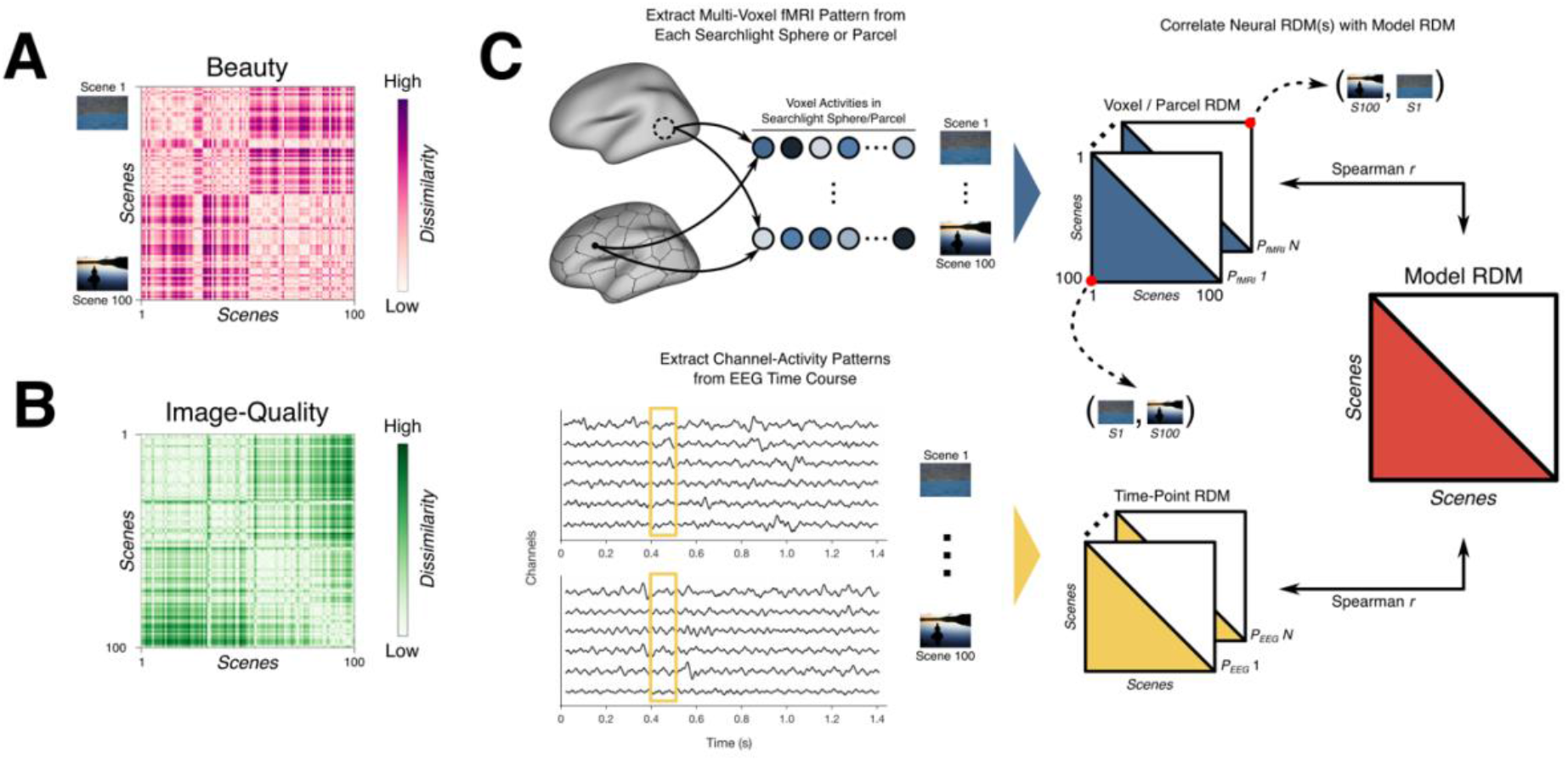
Illustration of RSA approach for fMRI and EEG. *A.* Beauty model RDM. We calculated the pairwise beauty-rating differences for all stimulus combinations using the behavioral data obtained in the fMRI and EEG studies. The beauty models from the fMRI and EEG studies were highly correlated (Spearman’s *r* = 0.88). We thus averaged the beauty predictors based on the ratings from the fMRI and EEG experiments, yielding the final beauty model RDM. *B.* Image-quality model RDM. We obtained image-quality ratings from a separate sample of *N* = 46 participants in an online survey. Analogous to the construction of the beauty model RDM, the image-quality model RDM consists of the pairwise image-quality rating differences for all stimulus combinations. Axes identical to *A*. *C*. Illustration of fMRI (top) and EEG (bottom) RSAs. We performed both a whole-brain searchlight RSA and a spatially discrete RSA in each parcel on the fMRI data. For the searchlight RSA, we constructed voxel-wise neural RDMs based on the average activity in a given searchlight sphere around a given voxel. To construct parcel-wise neural RDMs, we averaged the voxel-wise RDMs falling within the bounds of each cortical parcel, resulting in a single matrix for each parcel. Finally, the voxel- or parcel-wise RDMs were Spearman-correlated with the model RDM of interest. For the EEG RSA, we constructed time-point-wise neural RDMs based on the channel activities at each bin, which we then Spearman-correlated with the model RDM of interest.

#### Image-Quality Ratings

In order to model potential effects of image quality (i.e., how well or poorly a certain scene was captured by the photograph, independently of the content), we additionally collected image-quality ratings in an online survey completed by a separate group of 46 participants (*µ_age_* = 23.8 years, *SD_age_* = 4.0 years, 31 female). Survey participants were explicitly instructed that their ratings should reflect whether the scene images are good or bad photographs (in terms of color, contrast, saturation etc.) rather than scenes with more or less beautiful content. Each stimulus was rated on a scale from one (worst image quality) to seven (best image quality). Finally, we constructed an image-quality predictor RDM similarly to the beauty-predictor RDM: we first averaged the ratings across participants and then calculated the pairwise image-quality differences for all stimulus combinations (Figure 2B). The resulting image-quality RDM was only weakly correlated with the beauty-rating RDM (Spearman’s *r* = 0.12).

#### fMRI Searchlight and Parcel RSAs

To investigate whether visual, frontal, and DMN regions are involved in the representation of beauty in a spatially unbiased way, we performed a whole-brain searchlight RSA (Figure 2C, top). We defined a spherical searchlight of 300 voxels, and computed RDMs from patterns of beta values across the voxels falling within this sphere. These RDMs were then correlated to the beauty RDM using Fisher-transformed Spearman correlations. The searchlight RSA was performed in individual subjects. Finally, we computed a group-level searchlight map (*t*-test against zero, *p* < 0.001 voxel-wise, *p* < 0.05 FWE-corrected at the cluster level, *k* ≥ 100 voxels).

We performed an additional RSA by Spearman-correlating each brain parcel’s RDM with the beauty-rating RDM to investigate how strongly each spatially discrete cortical parcel represents beauty (Figure 2C, top). All correlations were Fisher-transformed and FDR-corrected for the number of tests (*t*-test against zero; *p_FDR_* < 0.05).

Both the searchlight and parcel RSA were repeated analogously with the exception that we now used partial Spearman correlations to additionally control for image quality by adding the image-quality rating RDM as a nuisance variable. This allowed us to examine in how far beauty judgments are influenced by the quality of the rated image. Finally, all thusly mentioned analyses were repeated with image quality as the main predictor. That is, we correlated the searchlight and parcel RDMs with the image-quality rating RDM while controlling for beauty ratings.

#### EEG RSA

We also performed RSA on the EEG data to investigate the representational dynamics of beauty judgments by correlating (Spearman correlation) each subject’s neural RDM from each time point with the beauty-rating RDM (Figure 2C, bottom). All correlations were Fisher-transformed and FDR-corrected for the number of tests. This yielded a representational time course of beauty judgments. Analogous to the fMRI RSA, this analysis was repeated using partial Spearman correlations while controlling for image quality, and with image quality as the main predictor, again controlling for beauty in a separate analysis.

#### EEG-fMRI Fusion

To combine the spatial resolution of the fMRI recordings and the temporal resolution of the EEG recordings, we conducted a model-based fusion analysis (Figure 3; Cichy & Oliva, 2020; Hebart et al., 2018). Our approach combined commonality analysis (Seibold & McPhee, 1979) with permutation testing to assess how much variance the beauty ratings explain in the shared EEG-fMRI representational similarity while preserving the original covariance structure of each predictor.

**Figure 3.**
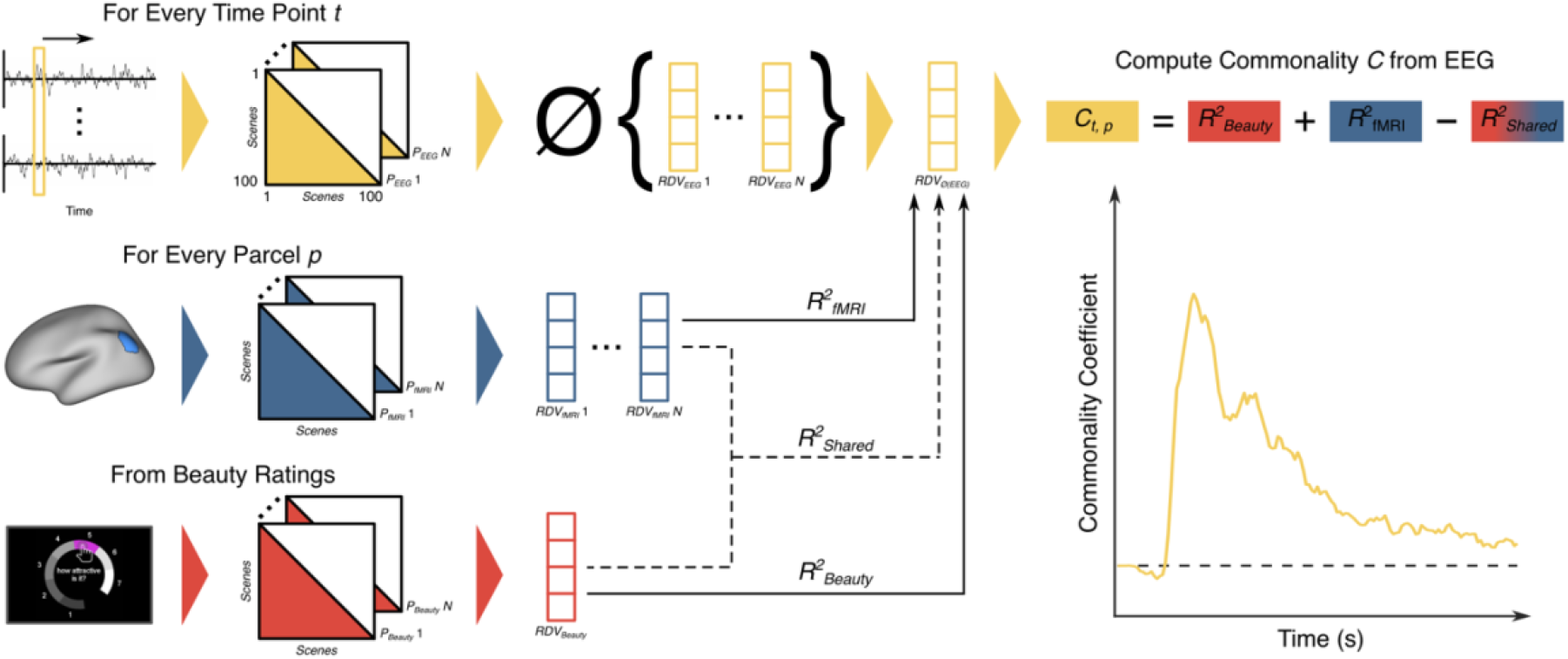
Fusion schematic. We first reshaped the lower triangle of each time point’s EEG RDM into an RDV and averaged the time-point-wise RDVs across participants. We also vectorized the parcel RDMs (without averaging across participants) and beauty-rating model RDM. All resulting RDVs were normalized to mean zero and a standard deviation of one before being entered into the commonality analysis. This involved calculating the variance explained by the beauty ratings in the EEG representational similarity (*R^2^* ) by regressing the EEG RDV from each time point onto the beauty RDV. We analogously computed the variance explained by each parcel’s representational similarity structure in the EEG (*R^2^* ), as well as the shared representational similarity between the beauty ratings and fMRI (*R^2^* ). The commonality coefficient *C* for each time point *t* and parcel *p* was then computed as the sum of *R^2^* and *R^2^* minus *R^2^* . Finally, we determined significant time points in each parcel’s commonality time course via a permutation test, and afterward FDR-corrected each time course for the number of time points (see Methods for details).

We first reshaped the lower triangle of each EEG participant’s RDM at every time point into a representational dissimilarity vector (RDV) and, for each bin, averaged the RDVs across participants. We also reshaped the RDMs of each brain parcel as well as the beauty and image-quality ratings into RDVs. Importantly, unlike the EEG data, the fMRI RDVs were not averaged across participants. Next, the EEG, fMRI, and rating RDVs were normalized (*µ* = 0, *SD* = 1) and entered into the commonality analysis. This was conducted by regressing the time-point-wise EEG RDVs onto the beauty-model RDV, yielding an index of the variance explained by the beauty ratings in the EEG data (*R^2^_Beauty_*) for each time point. This process was repeated for the RDVs of each fMRI parcel, resulting in an index of the variance shared between the representational similarities of the EEG data at a given time point and each brain parcel (*R^2^_fMRI_*). Finally, we also computed the shared variance between *R^2^_Beauty_* and *R^2^_fMRI_* (i.e., *R^2^_Shared_*), which allowed us to calculate the commonality coefficient *C* for each time point *t* and parcel *p* as:

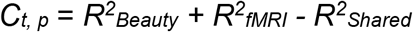

*C* is thus the shared variance in the representational similarities between EEG that is uniquely explained by the beauty ratings. We took the same approach for the additional fusion analysis in which we controlled for image quality, with the exception that all variables were residualized against the vectorized image-quality ratings via partial regression. In other words, we regressed each variable onto the image-quality ratings and performed the commonality analysis as described previously using the residuals of each of these regressions.

Given that collinearities between the predictors can artificially inflate commonality coefficients, we combined commonality analysis with a novel permutation-testing approach. Specifically, we sign-permuted both the beauty and parcel predictors 2,500 times and then fitted the original predictors to their shuffled counterparts, resulting in a strict lower bound for *C*. More precisely, this yielded two surrogate *R^2^* null distributions. We conservatively combined both distributions into one by retaining the maximum value for each permutation. For each time point and parcel, we then computed *p*-values by calculating the proportion of the permuted values of *C* that were smaller than the observed *C*. Finally, we corrected each parcel’s *p*-values for the number of time points using FDR-correction.

## Results

### Correlates of Scene Beauty Span a Large Network of Cortical Regions

We first performed a whole-brain searchlight RSA on the fMRI data. We used a spherical searchlight (300 voxels) centered on each voxel, from which we extracted beta values for each scene and constructed an RDM in which every entry was computed as one minus the correlation of beta patterns across the searchlight sphere for each pair of scenes. A model RDM was constructed from the pairwise rating differences, with each RDM entry reflecting the absolute difference in beauty ratings for two scenes. Finally, we computed Spearman correlations between the fMRI and model RDMs, yielding a whole-brain map (uncorr. *p* < 0.001 at the voxel level; *p_FWE_* < 0.05 at the cluster-level; *k* ≥ 100 voxels) indexing how well local activity patterns reflected behavioral scene-beauty ratings (Figure 2C, top).

This analysis revealed that beauty is represented across widespread cortical regions (Figure 4A). On the lateral brain surface, activation spanned ventral occipitotemporal cortex through superior parietal cortex, as well as lateral and dorsal prefrontal cortex (PFC), medial and inferior frontal gyrus, and the anterior insula. Sparse significant activity was additionally observed in anterior temporal regions, particularly the left middle temporal gyrus. On the medial surface, significant clusters emerged in precuneus, posterior cingulate cortex (PCC), dorsal and ventral aspects of PFCm, and OFC. Overall, the strongest effects were observed in precuneus, PCC, lateral occipitotemporal cortex (LOTC), and the inferior parietal lobule (IPL). In line with the initially discussed literature, the searchlight RSA revealed substantial overlap with the DMN, as evidenced by significant clusters in lateral parietal cortex, precuneus, PCC, and dorsal and ventral PFCm (cf. Raichle, 2015). Moreover, the extensive cluster in LOTC suggests the involvement of category-specific visual representations (Op de Beeck et al., 2019; Silson et al., 2016), such as scene-selective activity in the occipital place area and caudal IPL (Epstein & Baker, 2019).

**Figure 4.**
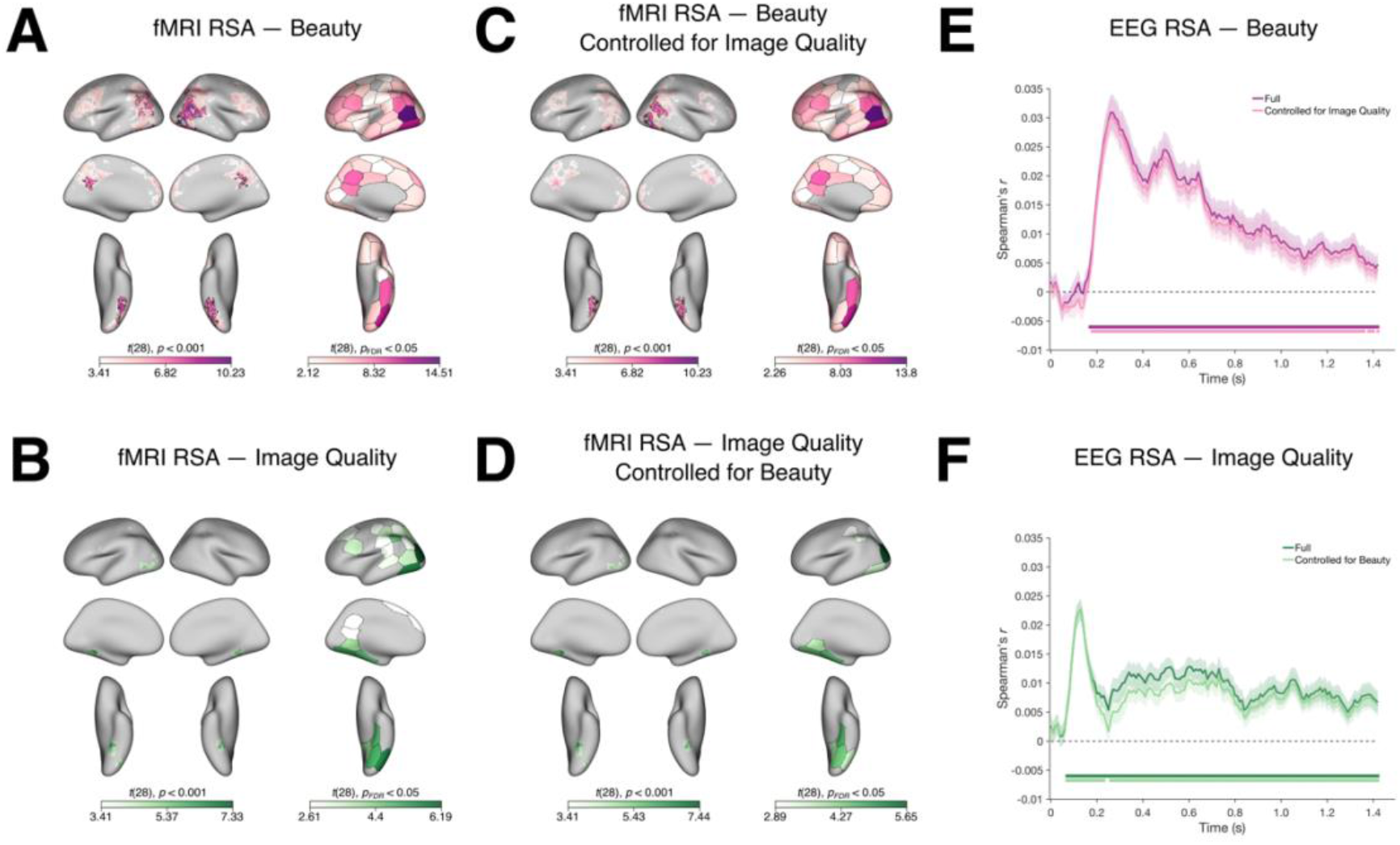
RSA results for fMRI and EEG. *A.* fMRI RSA reveals widespread beauty-related activations, particularly in high-level visual cortex and the DMN. Whole-brain searchlight map shown on the left (*t*-test against zero, voxel-wise uncorrected *p* < 0.001; areas outlined in black indicate significance at the cluster level, *pFWE* < 0.05, *k* ≥ 100) and parcel map on the right (Spearman correlation coefficient between parcel RDM and beauty model RDM, *t*-test against zero, FDR-corrected for the number of parcels). *B*. Image-quality representations have a distinct activation profile compared to beauty (structure and statistics identical to *A*.). *C.* Representations of beauty persist in a widespread cortical network after controlling for image quality (structure and statistics identical to *A.* and *B.*). *D.* Visual cortex represents image quality independently of perceived beauty (structure and statistics identical to *A.–C.*). *E.* Representations of beauty emerge early and are sustained over time, even when image quality is controlled for. To perform EEG RSA, we correlated each subject’s time-point-wise neural RDMs with the beauty model RDM. In the control analysis, we accounted for image quality via partial correlations. All correlations were Fisher-transformed. Markers indicate significant time points (*t*-test against zero, *p* < 0.05 FDR-corrected for the number of time points). *F.* Image-quality representations have qualitatively different temporal dynamics than beauty representations: they emerged within 100ms followed by sustained significant activity and persisted when accounting for the beauty ratings. Analyses performed analogously to *E*.

We additionally performed the same searchlight RSA while controlling for the image-quality ratings (Figure 4C), again yielding widespread representations of beauty, albeit at the absence of significant activity previously found in left LOTC and IPL as well as bilateral precuneus. Surprisingly, this suggests that beauty representations in predominantly visual areas (bilateral ventral and right lateral occipitotemporal cortices) are not explained by the lower-level visual features presumably captured by the image-quality ratings. Given their locus in high-level visual cortex, beauty representations are thus likely to be abstract, yet rooted in perceptual features.

In addition to the searchlight RSA, we also performed RSA in 50 *a priori* defined, bilateral brain parcels. Here, we averaged the RDMs of the voxels falling within both the left and right aspects of a given parcel and correlated the resulting RDM with the beauty ratings. This analysis revealed a similarly widespread representational profile that largely overlapped with the areas revealed by the initial searchlight RSA (Figure 4A). Indeed, controlling for image quality in the parcel RSA did not strongly affect the representational topography compared to the uncontrolled RSA (cf. Figures 4A, C). However, it is possible that a pattern of results similar to the whole-brain searchlight RSA was masked by the fact that the parcel RDMs were collapsed across the hemispheres. Taken together, our results nonetheless show a widespread network of regions involved in representing beauty during an explicit beauty-judgment task, notably including regions in visual cortex and the DMN.

We further inspected the similarities (Spearman correlations) between all parcel RDMs to investigate the degree of representational overlap between parcels (Supplementary Figure 1). The highest inter-regional similarities were observed between high-level cognitive networks, namely, the control, default-mode, and salience networks, and the visual networks VisCent and VisPeri. This suggests that representations in these areas contain similar information, and furthermore, that representations of beauty are instantiated in a common neural code in perceptual and cognitive brain regions.

### Early and Sustained Representations of Scene Beauty

To characterize neural representations of beauty over time, we performed RSA on EEG data reported in Kaiser (2022a) and Nara et al. (2026). We performed this analysis for 143 time bins of 10ms each, centered on 5ms to 1425ms post-stimulus. For each time bin, we constructed an RDM in which entries were computed as one minus the correlation of multi-electrode response patterns between pairs of scenes. The time-resolved RDMs were then correlated (Spearman correlation) with the model RDM derived from the pairwise beauty-rating differences, resulting in a representational time course of scene beauty (Figure 2C, bottom). As reported previously, we find that representations of beauty arise early and are sustained over time (Figure 4E). The correlations between neural and beauty RDMs peaked early at 265ms post-stimulus onset.

### Representations of Image Quality Are Distinct from Scene Beauty

Next, we investigated how neural representations of beauty compare to neural representations of image quality. To this end, we used the same fMRI- and EEG-RSA frameworks as outlined previously (Figure 2C), except that we correlated the neural (searchlight, parcel, or time-point) RDMs with the image-quality, rather than the beauty ratings. Our findings suggest that image-quality representations are distinct from those of beauty. Both the searchlight and parcel RSAs showed generally sparser activations for image quality relative to beauty, with the strongest activity found in early and ventral visual cortices, and weaker activity in frontal, temporoparietal, and posterior cingulate cortices (Figure 4B). Importantly, image-quality representations persisted only in visual areas after accounting for beauty (Figure 4D), suggesting more localized and predominantly perceptual coding.

This finding is corroborated by the EEG RSA: correlations between neural and image-quality RDMs yielded a qualitatively different time course than correlations in the beauty RSA (cf. Figures 4E, F), reaching significance before 100ms, and peaking at 135ms post-stimulus onset followed by sustained significant activity. The correlations between the neural and image-quality RDMs remained significant over time even when controlling for the beauty ratings. Indeed, there was virtually no drop in correlations after controlling for beauty relative to the original RSA during the first 200ms post-stimulus onset. Taken together, our results indicate that the neural signature of image quality emerges earlier than beauty, likely driven by lower-level visual features. More importantly, image-quality representations were distinct from those of beauty, indicating that the brain is able to both extract, and abstract away from the quality of the image when judging the aesthetic appeal of its content. This suggests that beauty in the world is processed differently than the beauty of the very depiction of the world.

### The Temporal Dynamics of Perceived Beauty Predominantly Map onto High-Level Vision

In order to provide a spatiotemporally resolved view of beauty-related neural representations, we performed EEG-fMRI fusion. We conducted our analyses in 50 *a priori* defined bilateral brain parcels obtained via resting-state fMRI (Yan et al., 2023). First, for each time point in the EEG, we reshaped the lower triangle of each subject’s neural RDM into a column vector and averaged the resulting RDVs across subjects, yielding a single RDM for every time point. Second, we extracted an RDM for each fMRI participant from every parcel. Finally, we determined the shared variance between the representational similarities of every parcel and time-point RDM via regression. We then combined *commonality analysis* (Hebart et al., 2018; Seibold & McPhee, 1979) and a novel collinearity-preserving permutation test to determine how much variance of the shared EEG-fMRI representational similarity is explained by the beauty ratings (see Methods and Figure 3 for details).

The commonality coefficients in three parcels in lateral occipital and ventral occipitotemporal cortices indicated a correspondence between temporally and spatially resolved representations (Figure 5A). Given that these parcels are located in high-level visual cortex, we conclude that high-level visual information is particularly critical for representations of scene beauty and their reflection in EEG signals. The commonality time courses of the resulting parcels follow highly similar trajectories, peaking between 250 and 270ms, followed by a secondary peak around 500ms and a gradual approach to baseline. This provides evidence that representations of beauty emerge in parallel across high-level visual regions, rather than following a temporal hierarchy. Although no other clusters yielded significant commonality scores, this parallel emergence was, qualitatively, observed across a larger network of brain parcels (see Supplementary Figure 2).

**Figure 5.**
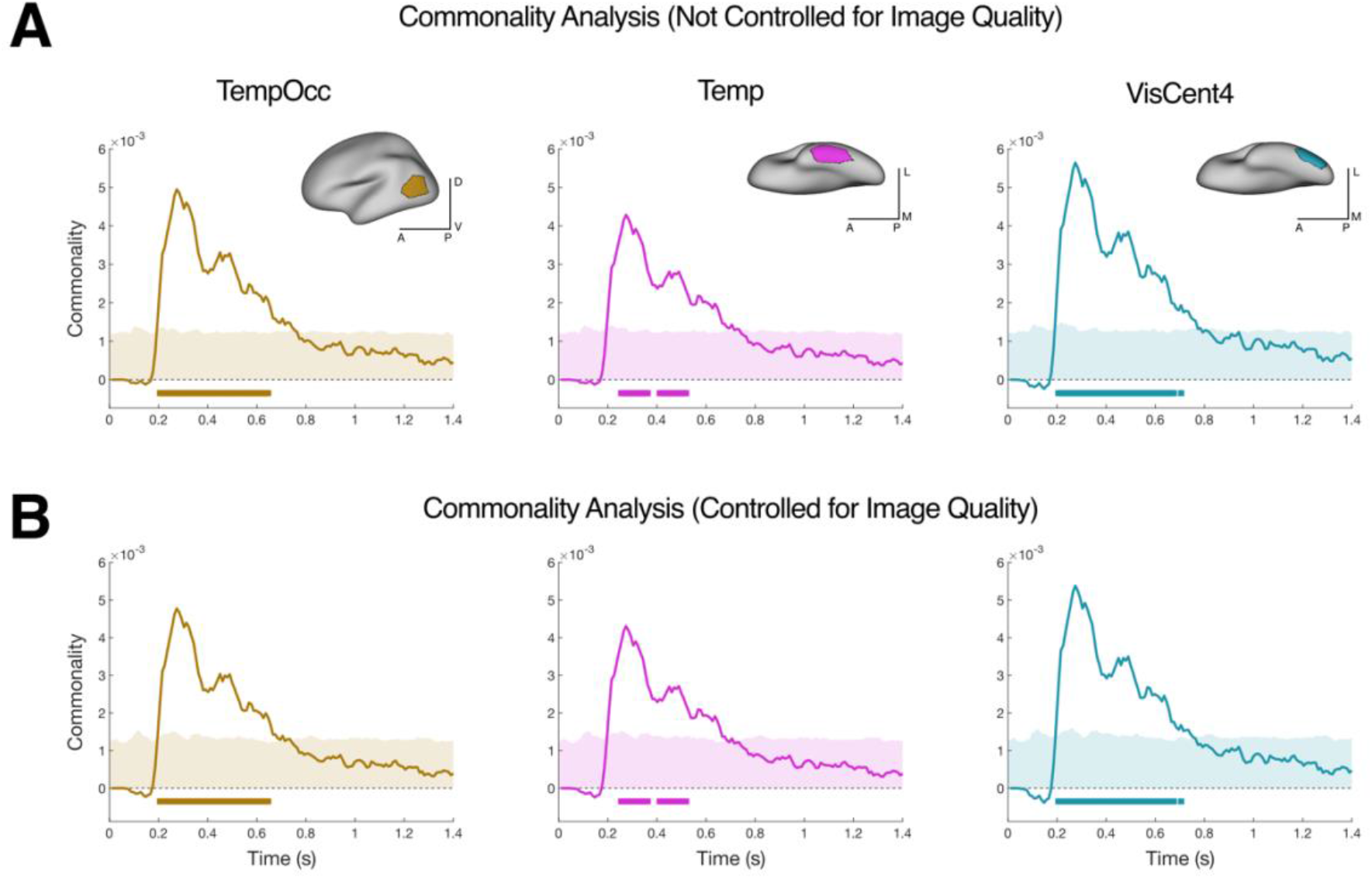
EEG-fMRI fusion results. *A.* Spatiotemporal representations of scene beauty map onto lateral occipitotemporal cortex (TempOcc) and ventral occipitotemporal cortex (Temp, VisCent4), emphasizing the importance of perceptual stimulus features for processing the perceived beauty of natural scenes. Shaded areas represent the 95^th^ percentile of commonality-score distributions obtained via permutation tests (see Methods for details). Markers underneath the line plots indicate significant time points (*t*-test against zero, *p* < 0.05 FDR-corrected for the number of time points). Abbreviations: D: dorsal, V: ventral, L: lateral, M: medial, A: anterior, P: posterior. *B.* Representations of scene beauty in high-level visual cortex persist when controlling for image quality, suggesting that the brain abstracts away from stimulus-level features when processing the beauty of natural stimuli. Statistical testing identical to *A*.

Next, we conducted the fusion analysis while additionally controlling for the image-quality ratings. This analysis yielded highly similar commonality time courses compared to the previous fusion analysis (cf. Figure 5A, B), providing further evidence for the distinctiveness of beauty and image-quality representations. In summary, the results of our EEG-fMRI fusion suggest that representations of beauty most prominently map onto brain regions in lateral and ventral occipitotemporal cortices. These regions likely contain visual representations of beauty that are independent of image quality, emphasizing the importance of complex perceptual features in processing and judging the aesthetic appeal of naturalistic stimuli.

## Discussion

In this study, we used RSA on EEG and fMRI data recorded while participants observed natural scene photographs to isolate neural representations of beauty and image quality in space and time, as well as model-based EEG-fMRI fusion to isolate the spatiotemporal dynamics of scene-beauty representations across the entire brain. Our RSA results confirm previous findings of early and sustained representations of beauty that span a wide network of brain regions. We also present evidence that the neural representations of image quality emerge automatically and are independent of representations of beauty, which is indicative of distinct processing modes for the perception of beauty in the outside world and in depictions of the world. Finally, the fusion analysis suggests that time-varying beauty representations predominantly map onto high-level visual areas, highlighting the importance of perceptual stimulus features in the processing of aesthetic appeal.

Early and sustained representations of beauty have been found previously (Kaiser, 2022a, 2022b; Kaiser & Nyga, 2020), even with varying stimulus-presentation times or task demands (Nara et al., 2026). Such early beauty-related representations may, in principle, conflate features of the world (the vividly green color of a tree) with features of the image (the blurriness of a bad photograph). Here, we show that that beauty and image quality are dissociable in the brain. First, we were able to recover neural signatures associated with the processing of image quality even though the neuroimaging participants had not been explicitly instructed to rate or pay attention to the quality of the stimuli. Second, we found that beauty-related representations had a different spatial and temporal profile than image-quality-related representations, which emerged earlier and mostly in visual cortex. Beauty and image quality may already be separated during visual processing, where image quality may be reflected in lower-level image statistics (e.g., spatial frequency, contrast) than beauty. When judging the beauty of what is depicted in a photograph, the brain may abstract away from such image-level features to distill the beauty of the scene that is depicted. While our results indicate independent processing of perceived quality and beauty, it is possible that explicit instructions to (additionally) rate image quality or more prominent quality degradations significantly affect representations of beauty (Tinio et al., 2011; Tinio & Leder, 2009). Future studies should further investigate the dependency of beauty in the world versus the quality of depictions of the world, for instance, by manipulating attention towards image quality or content, or by manipulating an image’s (artistic) style (Boger & Firestone, 2025).

Our fMRI analyses demonstrate that a wide range of functionally distinct brain areas are involved in the representation of beauty. However, the EEG-fMRI fusion seems to suggest that beauty representations are predominantly rooted in perceptual features, given that time-varying representations were localized to brain parcels in high-level visual cortex. This prominent role of visual analysis is in line with computational work demonstrating that human beauty ratings can successfully be predicted from visual feature coding alone (Conwell et al., 2025; Iigaya et al., 2021). Notably, a prominent involvement of high-level visual cortex in representations of beauty does not indicate that other brain systems are not important. Instead, our results suggest that the information the brain uses to construct aesthetic value first and foremost is stimulus-based. Prior work has demonstrated that early visual representations of aesthetic appeal are more readily predicted by low-level visual stimulus features and high-level features predict aesthetic appeal in late visual cortex, while the construction of aesthetic value is computed in PFCm by integrating across these hierarchical features (Iigaya et al., 2023). According to this view, the strong correspondence between time-varying beauty representations and regional representations in high-level visual cortex indicates that perceptual representations form the critical backbone for perceiving visual beauty.

What visual features relevant to scene beauty are represented in high-level visual cortex? Rather than encoding aesthetic appeal directly, ventral and lateral occipitotemporal cortex appears to represent visual features that have been shown to influence aesthetic judgments, such as symmetry, curvature, and scene composition or structure (Palmer et al., 2013). For one, activity in ventral and lateral occipitotemporal cortex has been shown to be modulated by symmetry, with higher degrees of (vertical) symmetry eliciting greater activation (Keefe et al., 2018; Van Meel et al., 2019). Similarly, parts of ventral visual cortex and the fusiform gyrus (Brodmann area 37) are characterized by a preference for curvature (Vartanian et al., 2024; Yue et al., 2020). In general, high-level visual cortex subserves naturalistic perception by representing global structural features of objects and scenes such as shape or composition (Ayzenberg & Behrmann, 2022; Bougou et al., 2024; Kaiser et al., 2020; Kravitz et al., 2011). Since representations in high-level visual cortex are more holistic, that is, less part-based and highly invariant to transformations (Op de Beeck et al., 2019), they may provide the perceptual basis upon which further downstream areas evaluate aesthetic appeal.

Beyond the flexible representation of aesthetically relevant visual features, high-level visual cortex is characterized by computational principles that make it particularly well suited as the perceptual backbone for the processing of aesthetic appeal. Such principles include the degree to which image elements can be integrated into a whole (Nara & Kaiser, 2024) or the sparseness of stimulus-evoked neural activity (Tang et al., 2025). These findings are well in line with processing fluency accounts of aesthetic preferences, which posit that easier—or: more fluent—neural processing predisposes greater aesthetic appeal (Graf & Landwehr, 2015; Reber et al., 2004). Relatedly, a recently proposed model (Walther, 2026) suggests that fluent categorization of visual inputs predisposes liking by reliably creating categorization-related micro-rewards. Our findings are consistent with this idea, given that the surviving parcels we obtained in the fusion analysis comprise category-selective areas. Taken together, the previous ideas and our current results consistently highlight the of role high-level visual cortex in representing aesthetic appeal, emphasizing the importance of perceptual stimulus features.

On a more cautious note, the representational correspondence between the EEG and fMRI data peaking in high-level visual cortex may be related to the limitations of EEG recordings. As our commonality analysis hinges on predicting the variance in the EEG signal, it is possible that beauty-related representations in early visual cortex could not be recovered due to qualitative differences between the representational structures of early EEG and fMRI responses (see: Cichy & Pantazis, 2017). This does not only apply to early visual activity but representations all over the brain: as revealed by our qualitative whole-brain fusion results, the temporal dynamics of all brain parcels are highly similar and resemble the temporal dynamics found in the EEG RSA (Figure 4E, Supplementary Figure 2). The limited sensitivity of EEG to recover subtle fluctuations in cortical processing dynamics (e.g., see: Iamshchinina et al., 2022; Proklova et al., 2019), particularly from unsuitable dipole orientations, different neuroimaging populations, and a relatively small stimulus set may have limited our propensity to uncover subtle variations in stimulus patterns generated in different parcels at different time points. On this view, the localization of time-varying beauty signals to high-level visual cortex primarily reflects a disproportionate contribution of visual cortex to the EEG signals. This view is also consistent with beauty-related EEG signals being relatively robust to stimulus presentation time and task demands (Nara et al., 2026). Subsequent studies may employ fusion using MEG and fMRI data recorded from a matched participant population to better resolve spatiotemporal beauty representations outside high-level visual cortex (Cichy & Oliva, 2020; Cichy & Pantazis, 2017).

Taken together, our results provide a spatiotemporally resolved view of neural representations of scene beauty. While judging beauty involves a wide range of brain areas, our work highlights the importance of perceptual features in grounding beauty representations. Our results moreover suggest that the processing of beauty is independent of image quality. In other words, our brains spontaneously separate the beauty of what is depicted from the quality of the depiction itself.

## Acknowledgments

We thank Martin Hebart and Philipp Schumann for help with the permutation testing routine for the EEG-fMRI fusion analysis. MR imaging for this study was performed at the Bender Institute of Neuroimaging (BION) at Justus Liebig University Giessen, Germany.

## Author Contributions

Conceptualization: P.F. and D.K. Data curation: P.F. Formal analysis: P.F. Funding acquisition: D.K. Investigation: P.F. Methodology: P.F. and D.K. Project administration: P.F. and D.K. Resources: D.K. Software: P.F. and R.S. Supervision: D.K. Validation: R.S. and D.K. Visualization: P.F. Writing—original draft: P.F. Writing— review and editing: R.S. and D.K.

## Availability of Data and Materials

All the stimuli, code, and data needed to evaluate the conclusions in the paper are present in the paper and/or the Supplementary Materials. Data, materials, and code can be found on Zenodo (single-subject beta maps; 10.5281/zenodo.16911734) and OSF (everything else; https://doi.org/10.17605/OSF.IO/S7ZBE ).

## Supplement

**Supplementary Figure 1.**
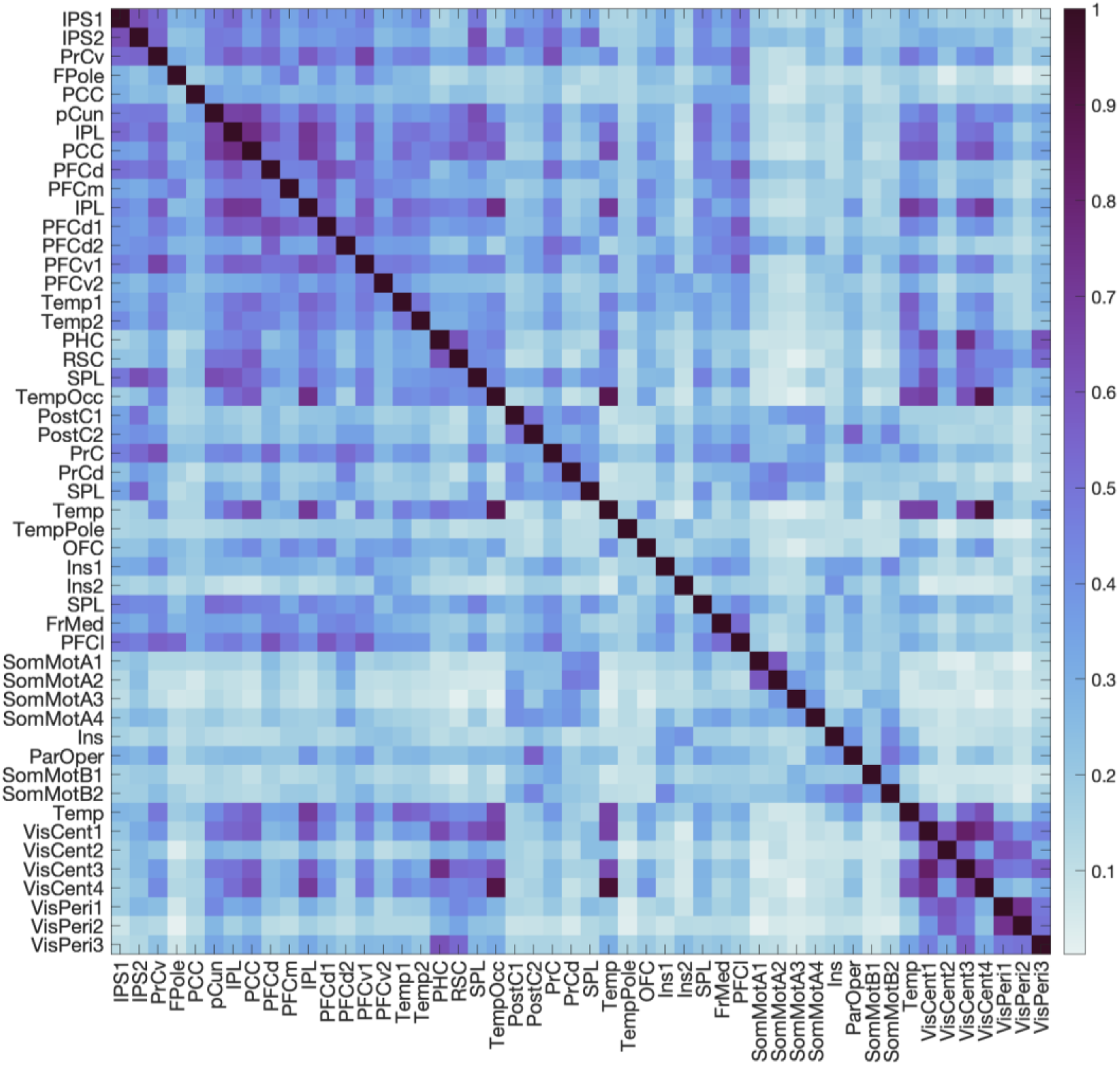
Correlations (Spearman) between all parcels’ representational dissimilarity structures. To investigate the amount of representational overlap between parcels, we vectorized each parcel’s RDM and calculated Spearman correlations between all combinations. The highest amount of overlap was observed within and between high-level cognitive networks (control: IPS, PrCv, PCC, pCun; DMN: IPL, PCC, PFC, Temp; dorsal attention: SPL, TempOcc; salience: SPL, FrMed, PFCl; temporoparietal: Temp) and visual networks (VisCent, particularly VisCent 4).

**Supplementary Figure 2.**
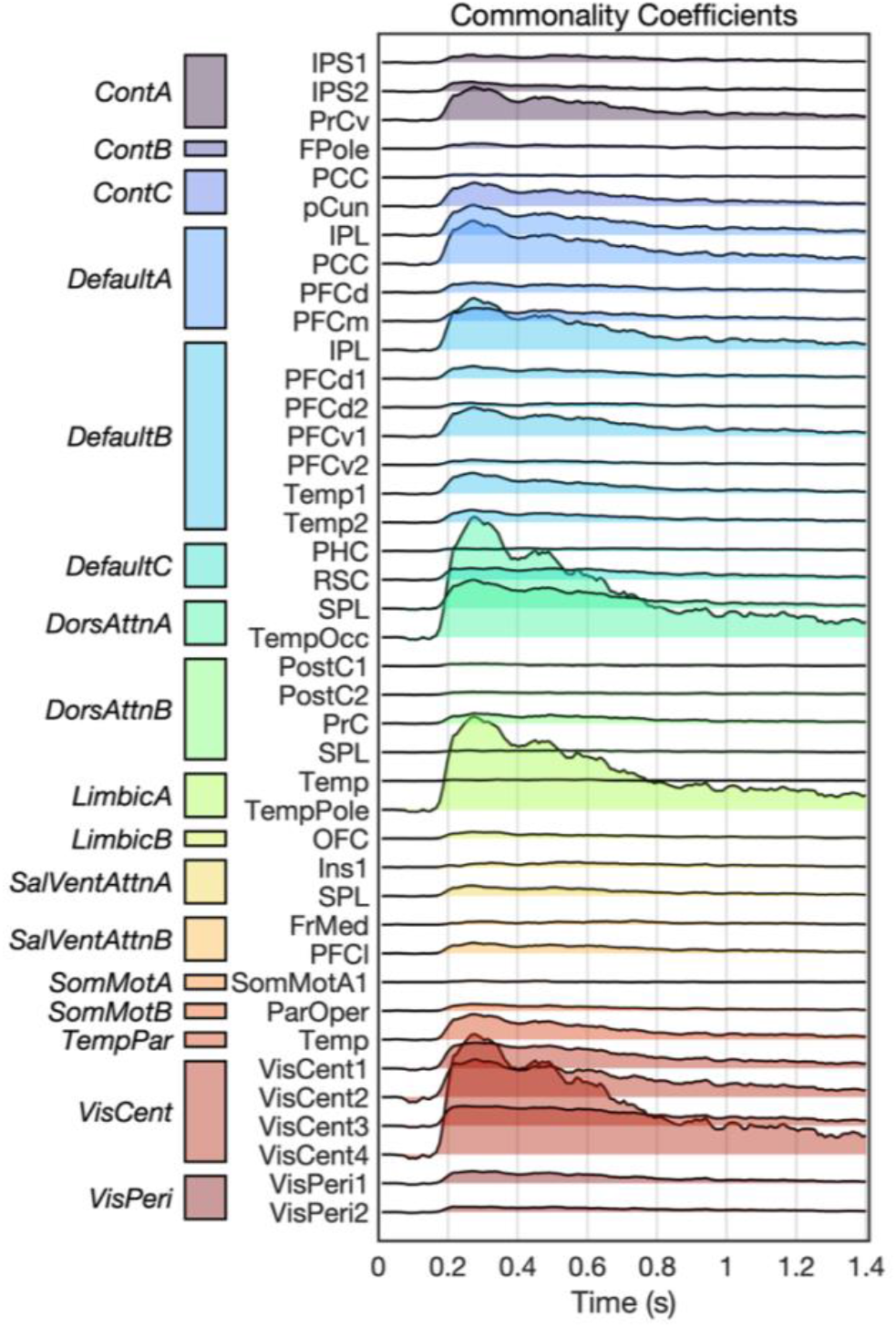
Qualitative results of the EEG-fMRI fusion analysis. We additionally conducted an uncorrected commonality analysis, and our results suggest that spatiotemporal representations of scene beauty emerge in parallel across widespread regions of cortex.

